# Optimal use of statistical methods to validate reference gene stability in longitudinal studies

**DOI:** 10.1101/545749

**Authors:** Venkat Krishnan Sundaram, Nirmal Kumar Sampathkumar, Charbel Massaad, Julien Grenier

## Abstract

Multiple statistical approaches have been proposed to validate reference genes in qPCR assays. However, conflicting results from these statistical methods pose a major hurdle in the choice of the best reference genes. Recent studies have proposed the use of a minimum of three different methods but there is no consensus on how to interpret conflicting results. Researchers resort to averaging the ranks or attributing a weighted rank to candidate genes. However, we report here that the suitability of these validation methods can be influenced by the experimental setting. Therefore, averaging the ranks can lead to suboptimal assessment of stable reference genes if the method used is not suitable for analysis. As the respective approaches of these statistical methods are different, a clear understanding of the fundamental assumptions and the parameters that influence the reference gene stability calculation is necessary. In this study, the stability of 10 candidate reference genes (*Actb, Gapdh, Tbp, Sdha, Pgk1, Ppia, Rpl13a, Hsp60, Mrpl10, Rps26*) was assessed using four common statistical approaches (GeNorm, NormFinder, Coefficient of Variation or CV analysis and Pairwise ΔCt method) in a longitudinal experimental setting. We used the development of the cerebellum and the spinal cord of mice as a model to assess the suitability of these statistical methods for reference gene validation. GeNorm and the Pairwise ΔCt were found to be ill suited due to a fundamental assumption in their stability calculations. Highly correlated genes were given better stability ranks despite significant overall variation. NormFinder fares better but the presence of highly variable genes influences the ranking of all genes because of the algorithm’s construct. CV analysis estimates overall variation, but it fails to consider variation across groups. We thus highlight the assumptions and potential pit-falls of each method using our longitudinal data. Based on our results, we have devised a workflow combining NormFinder, CV analysis along with visual representation of mRNA fold changes and one-way ANOVA for validating reference genes in longitudinal studies. This workflow proves to be more robust than any of these methods used individually.

**Additional Information:** Competing Interests – The authors declare no conflict of interest.

## Introduction

Relative expression of target genes in qPCR assays require accurate normalization of mRNA quantities using stably expressed internal standards also called reference genes (1). In theory, such a gene is presumed to be stably expressed across all groups and samples (2). This is seldom the case. There is no universal reference gene that fulfils this criterion and the choice of a good reference becomes highly subjective depending on several factors such as the sample/tissue type, experimental condition and sample integrity (3,4). Many statistical methods have been proposed to help researchers identify stable reference genes from a predetermined set of candidates. These statistical approaches determine the stability of these candidates based on a unique set of assumptions and calculations. Therefore, the predictions of these methods can vary rather significantly based on the method used and the experimental setting (5–7). This observation, to our knowledge, has been constantly neglected in recent studies that validate reference genes. However, to address this issue, researchers average the stability ranks across different methods and calculate an overall “geometric mean rank” (8,9). Some studies also attribute a weighted rank (10–14). This approach is rather questionable, as it does not consider the strengths and weaknesses of each method for a given experimental setting.

In this study, we tested the stability of 10 candidate reference genes during early postnatal development of the cerebellum and spinal cord in mice. This experimental setup proves to be a good longitudinal model. The cellular microenvironment is both complex and dynamic complicating the determination of stable reference genes (15–17). We first observed that the use of arbitrary reference genes gives highly variable profiles of the target gene depending on the reference chosen. Therefore, to identify stable reference genes, we used four different statistical approaches – GeNorm (18), NormFinder (19), Coefficient of Variation analysis (CV) (20) and Pairwise ΔCt method (21). However, the stability ranking also varied significantly depending on either the tissue in question or the method used.

Instead of averaging the ranks or attributing a weighted rank, we analysed the suitability of these methods to identify their respective drawbacks in a longitudinal setting. We find that the ranking of all methods tested except CV analysis are influenced by the presence of genes with high overall variation. Furthermore, GeNorm and the Pairwise ΔCt method rankings are influenced by the expression pattern of all genes making their ranking inter-dependent. On the other hand, NormFinder and CV analysis prove to be more robust, only when they are used complementarily but not individually. Hence, we devised an integrated approach by combining CV analysis, NormFinder and visual representation of mRNA fold changes across experimental groups. We believe that this method provides more accurate estimates of stable reference genes. In summation, our study highlights the importance of choosing the right set of statistical methods and proposes a sound workflow to validate reference genes in a longitudinal setting.

## Results

### Normalization with an arbitrary reference gene

To demonstrate the bias in results that arise by using a single arbitrary reference gene, we chose 3 candidate reference genes – *Actb, Gapdh* and *Mrpl10* to normalize Myelin Basic Protein (*Mbp)* mRNA expression levels in the cerebellum (Fig. 1a). *Mbp* levels showed a sudden increase by 35 folds at P10 when normalized to *Gapdh*, peaking at around 50 folds at P15 before coming down to 30 folds at P23. On the contrary, when normalized to *Mrpl10, Mbp* expression showed a rather linear relationship with time, gradually increasing from 12 to 41 folds between P10 and P23. When normalized to *Actb*, the *MBP* levels increase by 15 and 30 folds at P10 and P15 then shoot up to more than 90 folds at P23. However, the un-normalised profile of *Mbp* using P5 as a calibrator (Fig. 1b) shows an almost linear profile increasing from 18 to 56 folds between P10 and P23.

**Figure 1:**
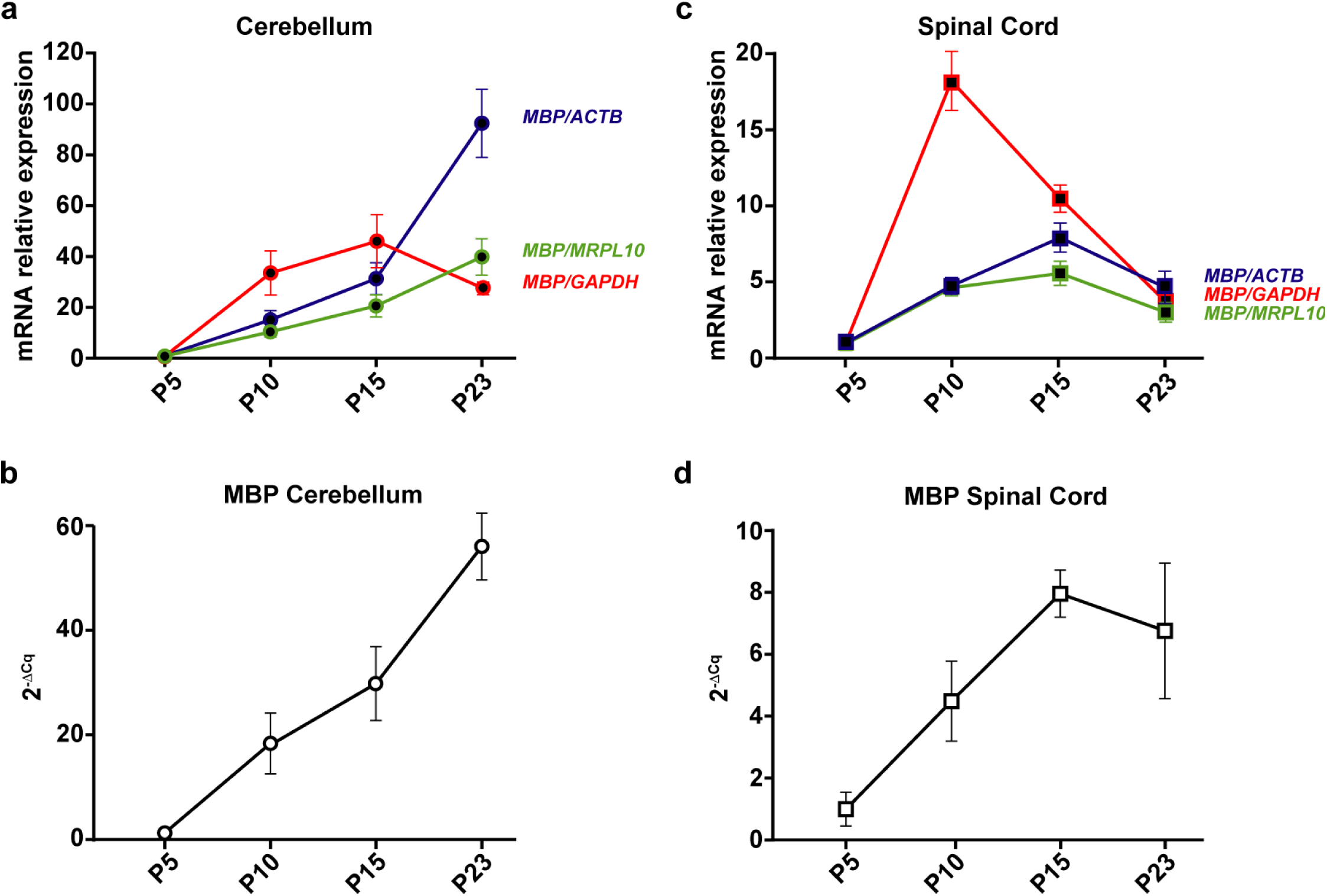
*Mbp* mRNA levels at post-natal day (P)5, 10, 15 and 23 in the cerebellum and spinal cord. P5 group is the experimental calibrator. (a) Cerebellar *Mbp* mRNA levels normalized using either *Actb, Mrpl10* or *Gapdh* mRNA quantities. (b) Un-normalized profile of cerebellar *Mbp* expressed as a fold change of mRNA quantities across groups (2-ΔCq). (c) Spinal cord *Mbp* mRNA levels normalized using either *Actb, Mrpl10* or *Gapdh* mRNA quantities. (d) Un-normalised profile of spinal cord *Mbp* expressed as a fold change of mRNA quantities across groups (2-ΔCq). Results are expressed as the Mean ± SD for each time point.

We observed similar contradictions in the spinal cord (Fig. 1c). *Mbp* levels normalized to *Gapdh* peak at P10 by 18 folds before reducing gradually to about 3 folds at P23. Normalizing with *Mrpl10* reveals a different kinetic where *Mbp* levels reach a plateau between P10 and P15 (around 4 and 6 folds respectively) before dropping down to 3 folds at P23. Whereas, normalizing with *Actb* reveals yet another profile where *Mbp* levels steadily increase from P5 to P15 in an almost linear fashion and decrease at P23 to around 5 folds. The un-normalised profile of *Mbp* (Fig. 1d) however remains linear till P15 and drops at P23.

### Raw expression profiles of candidate reference genes

Given the stark differences in *Mbp* expression profiles, we reasoned that the differences could be induced by intrinsic changes in the mRNA levels of candidate reference genes during development. To demonstrate this, we calculated the raw expression profiles of reference genes as fold changes of mRNA quantities across groups (Fig. 2&3). These intrinsic differences could indeed be shown as changes in Cq values for each gene across sample groups. However, Cq values are in the logarithmic scale and do not faithfully represent the magnitude of change in relative mRNA quantities that are always calculated as fold changes using a calibrator. Therefore, this data is better represented as fold changes.

**Figure 2:**
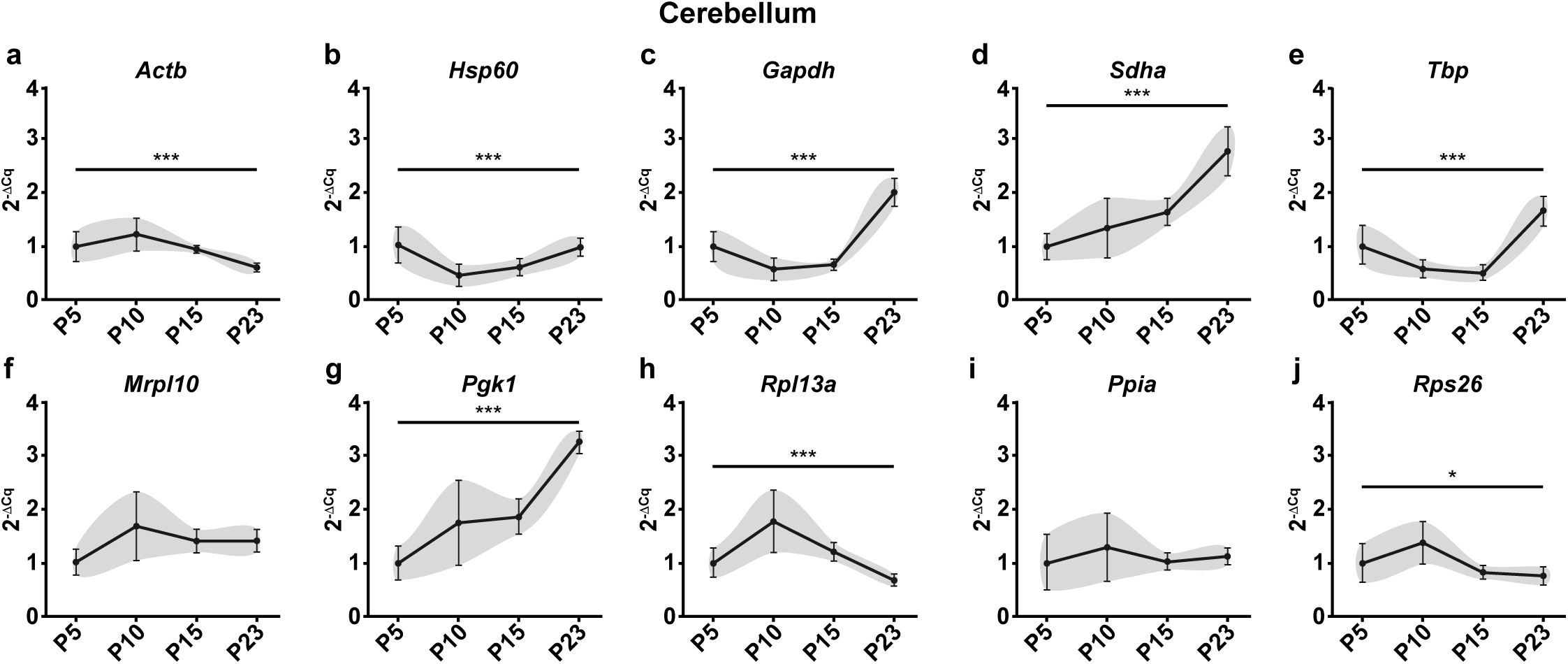
Raw expression profiles of reference genes expressed as fold changes across experimental groups (2^-ΔCq^) at P5, 10, 15 and 23 in the cerebellum. P5 group is the experimental calibrator. (a) *Actb*, (b) *Hsp60*, (c) *Gapdh*, (d) *Sdha*, (e) *Tbp*, (f) *Mrpl10*, (g) *Pgk1*, (h) *Rpl13a*, (i) *Ppia* and (j) *Rps26*. Results are expressed as the Mean ± SD for each time point. One-way ANOVA was performed to assess differences between the means of all groups. Statistical significance is denoted by P values: *P<0.05, **P<0.01, ***P<0.001. The grey area around each profile denotes the evolution of the intergroup and intragroup variation across time points.

**Figure 3:**
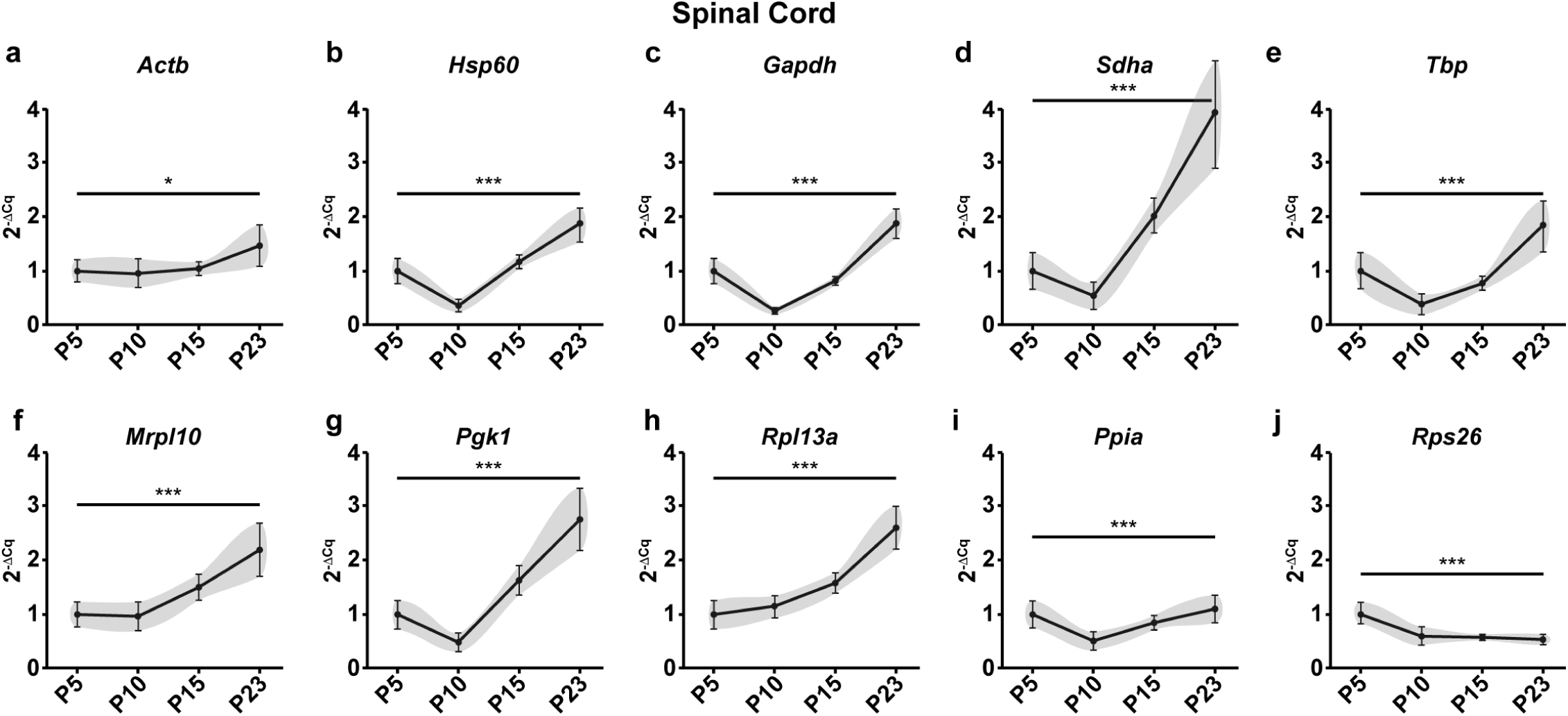
Raw expression profiles of reference genes expressed as fold changes across experimental groups (2^-ΔCq^) at P5, 10, 15 and 23 in the spinal cord. P5 group is the experimental calibrator. (a) *Actb*, (b) *Hsp60*, (c) *Gapdh*, (d) *Sdha*, (e) *Tbp*, (f) *Mrpl10*, (g) *Pgk1*, (h) *Rpl13a*, (i) *Ppia* and (j) *Rps26*. Results are expressed as the Mean ± SD for each time point. One-way ANOVA was performed to assess differences between the means of all groups. Statistical significance is denoted by P values: *P<0.05, **P<0.01, ***P<0.001. The grey area around each profile denotes the evolution of the intergroup and intragroup variation across time points.

The first statistical test to assess stability after visually representing the data was One-way ANOVA to determine if the mean mRNA levels across groups are significantly different from one another. In the cerebellum, 8 of the 10 reference genes tested (*Actb, Hsp60, Gapdh, Sdha, Tbp, Pgk1, Rpl13a* and *Rps26)* showed significant variation in the mRNA levels across time points (Fig. 2) Only 2 genes (*Mrpl10, Ppia)* showed no significant change. In the spinal cord, all tested genes showed significant variation in mRNA levels across the four time points (Fig 3).

These results taken together with the raw expression profiles of *Mbp* (Fig. 1b & 1d) show that intrinsic changes in mRNA levels of reference genes can indeed skew the normalized profile of *Mbp*. As a result, it causes a significant bias in the results and interpretations that ensue highlighting the importance of validating reference gene stability in longitudinal studies.

### Assessment of expression stability using multiple statistical approaches

The expression stability of candidate reference genes was analysed in both tissues using four well known statistical methods - CV analysis (20), NormFinder (19), GeNorm (18) and the Pairwise ΔCt method (21) (Table 2).

**Table 1:**
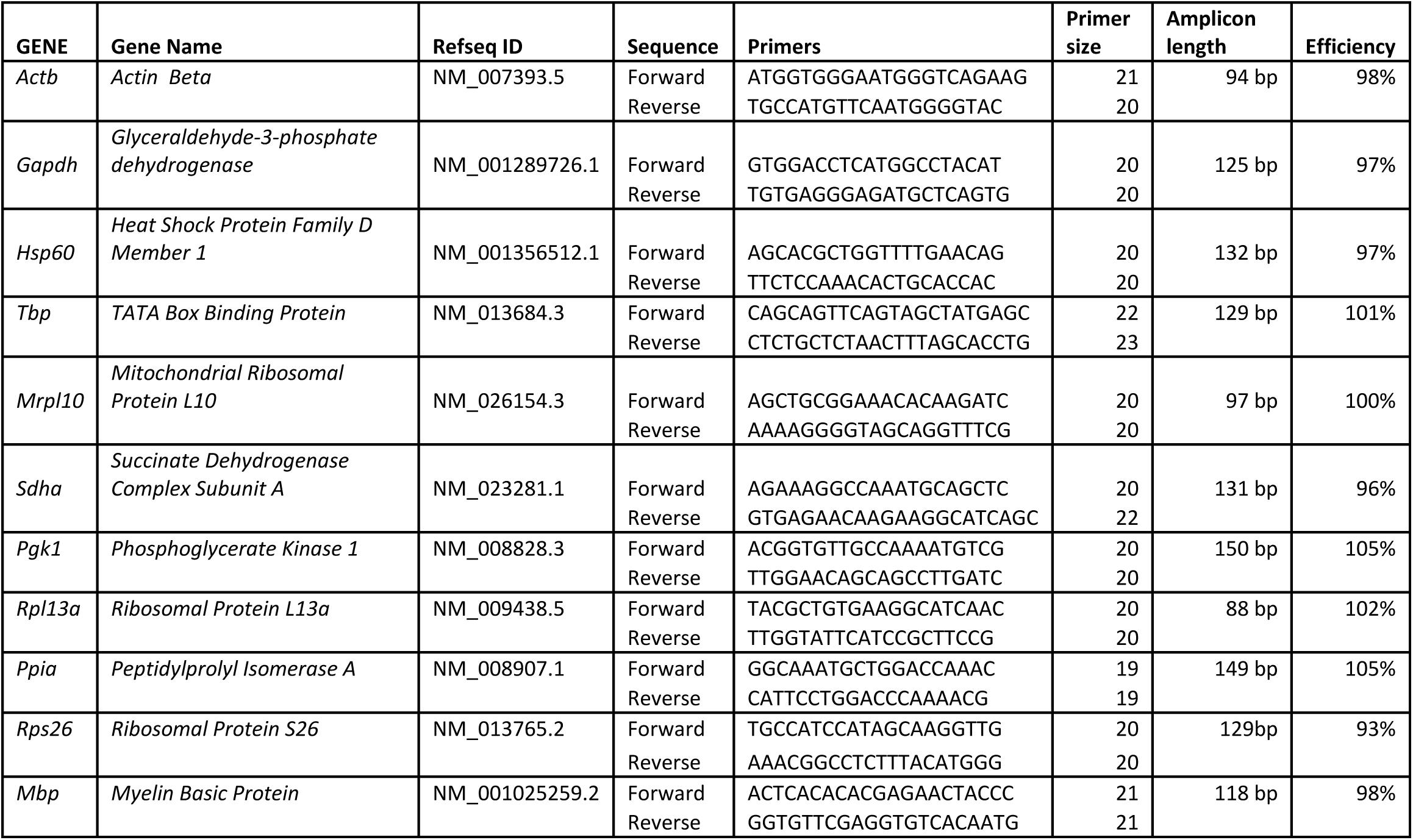
Details of the primer sequences used in the study.

**Table 2:** Expression stability of candidate reference genes in cerebellum and spinal cord evaluated using Coefficient of Variation (CV) Analysis, NormFinder, GeNorm and the Pairwise ΔCt method. Coefficient of Variation (CV) analysis is a simple descriptive statistical method where the Cq values of all candidate reference genes across samples are first linearized (2^-Cq^). Next the CV for each gene across all samples is calculated and expressed as a percentage. NormFinder uses a model-based approach where it calculates the stability of reference genes based on two parameters – the intergroup variation and the intragroup variation. The stability score denoted by the S value is a weighted measure of these two parameters. The most stable reference gene has the least S value. GeNorm attributes a stability score (M value), which is assessed by pairwise variation. The algorithm functions by first identifying two genes with the highest expression agreement and therefore high stability. It then calculates the expression variation of every other gene sequentially with respect to the previous genes chosen. Therefore, the ranking of the GeNorm algorithm always has 2 genes at the top with the same M value followed by other genes with higher M values indicating lower stability. The pairwise ΔCt method has the same rationale as the GeNorm method but evaluates the stability score differently. This approach calculates the differences in Cq values between one gene and the others across all samples. This is followed by the calculation of the standard deviation of these differences. A low Mean SD value indicates high stability. Refer to Supplementary Information for more details.

Cerebellum
| CV Analysis |  |  |
| --- | --- | --- |
| Gene | CV% | Rank |
| *Mrpl10* | 30.4 | 1 |
| *Actb* | 31.9 | 2 |
| *Rps26* | 35.3 | 3 |
| *Ppia* | 36.8 | 4 |
| *Hsp60* | 42.6 | 5 |
| *Rpl13a* | 44.7 | 6 |
| *Sdha* | 45.2 | 7 |
| *Pgk1* | 47.3 | 8 |
| *Tbp* | 57.2 | 9 |
| *Gapdh* | 59.1 | 10 |

| Norm Finder |  |  |
| --- | --- | --- |
| Gene | Stability S | Rank |
| *Ppia* | 0.31 | 1 |
| *Mrpl10* | 0.33 | 2 |
| *Sdha* | 0.47 | 3 |
| *Rps26* | 0.48 | 4 |
| *Pgk1* | 0.49 | 5 |
| *Hsp60* | 0.5 | 6 |
| *Actb* | 0.55 | 7 |
| *Rpl13a* | 0.58 | 8 |
| *Gapdh* | 0.62 | 9 |
| *Tbp* | 0.68 | 10 |

| GeNorm |  |  |
| --- | --- | --- |
| Gene | Stability M | Rank |
| *Gapdh* | 0.37 | 1 |
| *Tbp* | 0.37 | 1 |
| *Hsp60* | 0.44 | 2 |
| *Sdha* | 0.55 | 3 |
| *Pgk1* | 0.59 | 4 |
| *Mrpl10* | 0.63 | 5 |
| *Ppia* | 0.65 | 6 |
| *Rps26* | 0.69 | 7 |
| *Actb* | 0.71 | 8 |
| *Rpl13a* | 0.74 | 9 |

| Pairwise Delta C method |  |  |
| --- | --- | --- |
| Gene | Average SD | Rank |
| *Mrpl10* | 0.608 | 1 |
| *Ppia* | 0.652 | 2 |
| *Sdha* | 0.698 | 3 |
| *Rps26* | 0.699 | 4 |
| *Hsp60* | 0.745 | 5 |
| *Pgk1* | 0.751 | 6 |
| *Actb* | 0.758 | 7 |
| *Gapdh* | 0.813 | 8 |
| *Tbp* | 0.82 | 9 |
| *Rpl13a* | 0.83 | 10 |

Spinal Cord
| CV analysis |  |  | Norm Finder |  |  |
| --- | --- | --- | --- | --- | --- |
| Gene | CV% | Rank | Gene | Stability S | Rank |
| *Actb* | 28.10 | 1 | *Mrpl10* | 0.29 | 1 |
| *Ppia* | 32.80 | 2 | *Ppia* | 0.30 | 2 |
| *Rps26* | 36.00 | 3 | *Hsp60* | 0.30 | 3 |
| *Mrpl10* | 40.50 | 4 | *Rpl13a* | 0.35 | 4 |
| *Rpl13a* | 42.80 | 5 | *Tbp* | 0.39 | 5 |
| *Hsp60* | 49.40 | 6 | *Actb* | 0.44 | 6 |
| *Pgk1* | 57.70 | 7 | *Pgk1* | 0.45 | 7 |
| *Tbp* | 58.60 | 8 | *Gapdh* | 0.50 | 8 |
| *Gapdh* | 59.20 | 9 | *Sdha* | 0.62 | 9 |
| *Sdha* | 72.10 | 10 | *Rps26* | 0.83 | 10 |

| GeNorm |  |  | Pairwise Delta C method |  |  |
| --- | --- | --- | --- | --- | --- |
| Gene | Stability M | Rank | Gene | Average SD | Rank |
| *Mrpl10* | 0.19 | 1 | *Hsp60* | 0.535 | 1 |
| *Rpl13a* | 0.19 | 1 | *Mrpl10* | 0.536 | 2 |
| *Pgk1* | 0.39 | 2 | *Pgk1* | 0.544 | 3 |
| *Hsp60* | 0.44 | 3 | *Ppia* | 0.554 | 4 |
| *Tbp* | 0.46 | 4 | *Tbp* | 0.565 | 5 |
| *Ppia* | 0.48 | 5 | *Rpl13a* | 0.593 | 6 |
| *Gapdh* | 0.51 | 6 | *Actb* | 0.623 | 7 |
| *Sdha* | 0.53 | 7 | *Gapdh* | 0.665 | 8 |
| *Actb* | 0.55 | 8 | *Sdha* | 0.706 | 9 |
| *Rps26* | 0.63 | 9 | *Rps26* | 0.934 | 10 |

#### CV analysis

The CV analysis estimates the variation in the linearized Cq values (2^-Cq^) of a reference gene across all samples taken together (Supplementary Table S1). Therefore, lower CV would mean higher stability. CV analysis on the cerebellum samples revealed *Mrpl10, Actb and Rps26* as the top three stable reference genes. In the spinal cord, the top three reference genes are *Actb, Ppia, Rps26*.

#### NormFinder analysis

We next analysed our data using NormFinder which calculates a stability score (S) based on the intergroup and intragroup variations. In the cerebellum, NormFinder identified *Ppia, Mrpl10* and *Sdha* as the top three stable genes. In the spinal cord, the algorithm identified *Mrpl10, Ppia* and *Hsp60* as the top three stable reference genes.

#### GeNorm analysis

GeNorm calculates stability based on pairwise variation. The rationale is that if two genes vary similarly across all samples, then they are the most stable reference genes for that dataset. All other genes are ranked on their similarity to the expression of the top two genes. In the cerebellum, the GeNorm algorithm identified *Gapdh* and *Tbp* to be the most stable reference genes followed by *Hsp60*. This is in stark contrast to the genes identified by the previous two methods. In the spinal cord, the most stable reference genes were identified as *Mrpl10* and *Rpl13a* followed by *Pgk1*.

#### Pairwise ΔCt analysis

This method works on the same rationale as GeNorm but calculates the stability value (Mean SD) differently. It is calculated as the average standard deviation of the Cq value differences that the gene exhibits with other genes (Supplementary Table S2). In the cerebellum, Pairwise ΔCt analysis identified *Mrpl10, Ppia* and *Sdha* as the top three most stable reference genes. The ranking of this method and NormFinder are quite similar in the cerebellum. However, in the spinal cord the top three genes were identified as *Hsp60, Mrpl10* and *Pgk1*, noticeably different from the NormFinder rankings.

The overall ranking of genes across all methods recapitulated in Table 2 show that the stability ranking of reference genes can indeed vary significantly depending on the tissue studied and the method used for validation. This makes the identification of the best reference genes very cumbersome.

### Suitability of validation methods for longitudinal studies

We next analysed the suitability of these methods in a longitudinal setting highlighting the assumptions of each method and the factors that influence the stability scores.

#### CV analysis

As mentioned, the CV analysis adopts a direct approach by calculating the variance of a gene across all samples taken together. However, this method does not consider the variation across different time points or groups. For example, *Actb* is ranked 2^nd^ in the cerebellum (Table 2) as it exhibits low overall variation but the profile of the gene (Fig. 2a) tells us that it does vary across time points. This is a major concern in a longitudinal study as this method determines variation in just one dimension whereas the dataset exists in two dimensions.

#### NormFinder analysis

The NormFinder approach is more robust as it calculates the stability based on the intergroup and intragroup variation. However, on inspecting the algorithm further, we found that including genes with high overall variation can affect the stability rankings of all the genes. Consider the overall variation of *Actb* in the spinal cord (Fig. 3a). It has a CV of 28.1% (Table 2), which is the least among all genes in that group. Furthermore, the profile of *Actb* in the spinal cord is almost flat with minimal variation across groups (Fig 3a). Nevertheless, it is ranked 6^th^ in the NormFinder method. *Hsp60* and *Rpl13a*, on the other hand, with a higher overall variance and significant intergroup variation when compared to *Actb* (Fig. 3a, b and h) are ranked 3^rd^ and 4^th^. *Hsp60* is in fact attributed the same stability value as *PPIA* which is ranked 2^nd^. This possible flaw in the algorithm is because of the presence of genes with high overall variations. Hence, the NormFinder algorithm can potentially be improved after identifying and removing genes with high overall variation *(See Towards an integrated approach)*.

#### GeNorm analysis

The GeNorm algorithm as explained earlier ranks genes based on pairwise correlation. Hence, there is always a possibility that this method could estimate correlated and co-regulated genes to be highly stable. To validate the rankings of this algorithm we performed Pearson’s correlation on the linearized Cq values of all genes and samples (Fig. 4a & 4b, Supplementary Table S3 & S4 for the correlation matrices). We observed two distinct patterns of correlation in the cerebellum and the spinal cord, the cerebellum being more heterogeneous. In the spinal cord, almost all genes tested exhibit a positive correlation except *Actb* which correlates less and *Rps26* which exhibits negative correlation with most of the other genes. This is also evident from the raw expression profiles where we see that the raw expression profiles in the cerebellum are heterogeneous (Fig. 2). In the spinal cord, however, all the reference genes seem to have a similar profile except for *Actb* and *Rps26* (Fig. 3).

**Figure 4:**
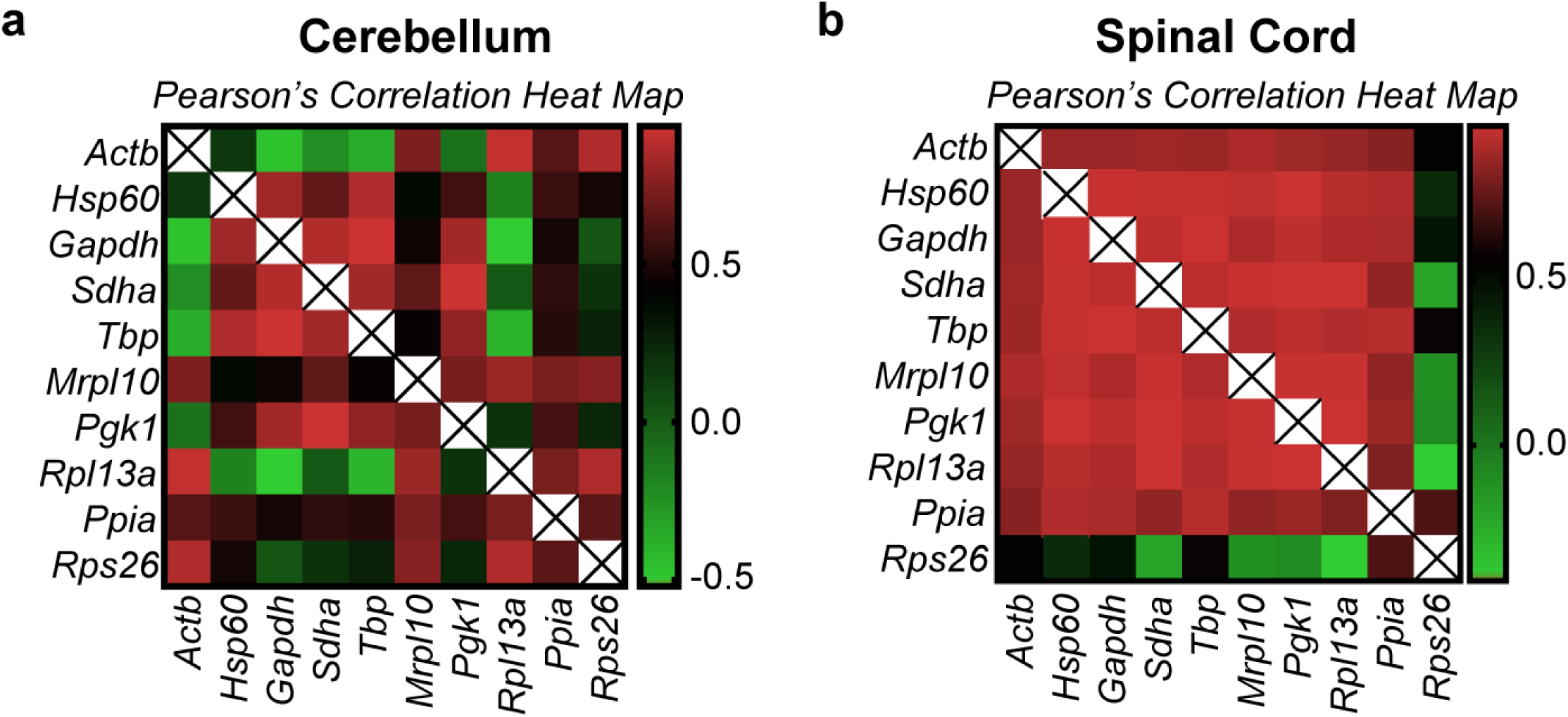
Pearson’s correlation heatmap of the linearized Cq values (2^-Cq^) of all genes. Pearson’s correlation of all genes was performed to identify the extent of expression correlation in (a) cerebellum and (b) spinal cord. The colour scheme is denoted next to each heatmap. The numerical values used in the colour scheme represent the Pearson’s r score. r = 0 (no correlation), r = +0.5 (positive correlation), r = −0.5 (negative correlation). r values closer to +1 or −1 denote strong positive and negative correlations respectively. The diagonal lines marked with an “X” are indicative of the correlation of a gene with itself and therefore is omitted from analysis.

Interestingly, the highest correlation in the cerebellum dataset was observed in *Gapdh*/*Tbp* with a Pearson’s r score of 0.937. This gene pair is ranked first in the GeNorm analysis (Table 2) but have the highest overall variation in the group (CV=59.1% and 57.2%). In the spinal cord, however, the top two genes according to GeNorm are *Mrpl10*/*Rpl13a* while the highest correlation is observed in *Sdha*/*Pgk1* with a Pearson’s r score of 0.97. Intriguingly, the genes that are classed below - *Pgk1, Hsp60* and *Tbp* have CV values of 57.8%, 49.4% and 58.6%, which are among the highest in the spinal cord group. This is because of their expression agreement with the top two genes. Seemingly stable genes such as *Ppia* (CV=32.8%, Fig. 3i) and *Actb* (CV=28.1%, Fig. 3a) are pushed to the lower half of the ranking (Table 2).

These results suggest that GeNorm tends to favour highly correlated genes in a heterogeneously correlated set of genes (cerebellum dataset). In a less heterogenous dataset (spinal cord) the ranking does not concur with the overall stability of the genes across samples assessed by visual representation (Fig. 3) and CV analysis (Table 2). These results taken together show that GeNorm’s ranking can be influenced by the extent of correlation in the dataset. This exclusion of possibly stable genes by GeNorm because of negative or less correlation was indeed foreseen by Anderson and colleagues when they proposed the NormFinder model. We find that to be true in our analysis.

#### Pairwise ΔCt Analysis

The pairwise ΔCt approach seems to work better in a heterogeneously correlated set of genes. The rankings of NormFinder and the pairwise ΔCt method are rather identical in the cerebellum. However, in the spinal cord apart from *Mrpl10* the other top genes - *Pgk1* and *Hsp60* - show high variation across samples as explained earlier. The limitations of this method can also be observed in the spinal cord. *Actb*, which shows a rather flat profile (Fig. 3a) is very distinct when compared to the others. It correlated very little with other genes (Fig. 4b, Supplementary Table S4). This increases the difference in the Cq values between *Actb* and others resulting in a high Mean SD (stability value) thereby attributing it with a lower rank. Therefore, although a gene varies very little across samples and groups, the pairwise ΔCt approach might attribute it with a lower rank if the profiles of the other genes are different. The same argument is also true for *Rps26* in the spinal cord (Fig. 3j, 4b)

In conclusion, we find that the ranking of all the methods except CV analysis is influenced by the inclusion of genes with high overall variation. Additionally, the pairwise expression stability methods (GeNorm and Pairwise ΔCt) could produce misleading results depending on the profile of the highly variable genes together with the extent of expression correlation between all the genes.

### Towards an integrated approach

The first step towards an integrated approach was to remove genes with high overall variance. CV analysis calculates the overall variance of each gene and it is the only method where the stability ranking of a gene is not influenced by others. We therefore used this method to objectively identify genes with high overall variance. We defined a threshold of CV = 50%. Genes that exhibit a CV above this value are taken to be highly variable and are excluded from further analysis. In the cerebellum, *Gapdh* and *Tbp* were removed and in the spinal cord *Gapdh, Tbp, Pgk1* and *Sdha* were removed from analysis. We next determined how this exclusion impacts the stability ranking of NormFinder, GeNorm and the Pairwise ΔCt approach in both tissues (Table 3).

**Table 3:** Revised rankings of candidate reference genes in cerebellum and spinal cord using NormFinder, GeNorm and the Pairwise ΔCt method. In the cerebellum, *Gapdh* and *Tbp* were excluded from analysis. In the spinal cord, 4 genes – *Sdha, Gapdh, Tbp* and *Pgk1* were excluded from analysis

Cerebellum
| Norm Finder |  |  |
| --- | --- | --- |
| Gene | Stability<br>S | Rank |
| *Mrpl10* | 0.19 | 1 |
| *Ppia* | 0.21 | 2 |
| *Rps26* | 0.45 | 3 |
| *Actb* | 0.45 | 4 |
| *Rpl13a* | 0.47 | 5 |
| *Sdha* | 0.51 | 6 |
| *Pgk1* | 0.52 | 7 |
| *Hsp60* | 0.64 | 8 |

| GeNorm |  |  |
| --- | --- | --- |
| Gene | Stability<br>M | Rank |
| *Actb* | 0.30 | 1 |
| *Rpl13a* | 0.30 | 1 |
| *Rps26* | 0.33 | 2 |
| *Mrpl10* | 0.40 | 3 |
| *Ppia* | 0.46 | 4 |
| *Sdha* | 0.57 | 5 |
| *Pgk1* | 0.62 | 6 |
| *Hsp60* | 0.67 | 7 |

| Pairwise Delta C |  |  |
| --- | --- | --- |
| Gene | Average<br>SD | Rank |
| *Mrpl10* | 0.529 | 1 |
| *Ppia* | 0.598 | 2 |
| *Rps26* | 0.614 | 3 |
| *Actb* | 0.65 | 4 |
| *Rpl13a* | 0.703 | 5 |
| *Sdha* | 0.714 | 6 |
| *Pgk1* | 0.756 | 7 |
| *Hsp60* | 0.823 | 8 |

Spinal Cord
| Norm Finder |  |  |
| --- | --- | --- |
| Gene | Stability<br>S | Rank |
| *Ppia* | 0.22 | 1 |
| *Mrpl10* | 0.25 | 2 |
| *Actb* | 0.28 | 3 |
| *Rpl13a* | 0.35 | 4 |
| *Hsp60* | 0.48 | 5 |
| *Rps26* | 0.64 | 6 |

| GeNorm |  |  |
| --- | --- | --- |
| Gene | Stability<br>M | Rank |
| *Mrpl10* | 0.19 | 1 |
| *Rpl13a* | 0.19 | 1 |
| *Actb* | 0.32 | 2 |
| *Ppia* | 0.39 | 3 |
| *Hsp60* | 0.47 | 4 |
| *Rps26* | 0.58 | 5 |

| Pairwise Delta C |  |  |
| --- | --- | --- |
| Gene | Average<br>SD | Rank |
| *Mrpl10* | 0.472 | 1 |
| *Ppia* | 0.49 | 2 |
| *Actb* | 0.494 | 3 |
| *Rpl13a* | 0.537 | 4 |
| *Hsp60* | 0.68 | 5 |
| *Rps26* | 0.78 | 6 |

#### Ranking changes in NormFinder

As expected, exclusion of highly variable genes changed the overall ranking of the NormFinder algorithm in the cerebellum (Table 3). Although the top two genes remained the same, the ranking of other genes changed. *Sdha, Hsp60* and *Pgk1* that were ranked higher than *Actb* (Table 2) now obtained ranks lower than *Actb* (Table 3). The top three genes after exclusion of *Gapdh* and *Tbp* were identified as *Mrpl10, Ppia* and the third position is shared between *Rps26* and *Actb*. The most stable genes identified by NormFinder after exclusion (Table 3) concurs with the CV analysis (Table 2) and with the expression profiles of the genes (Fig. 2a, 2f, 2i & 2j) wherein the most stable genes exhibit minimal variation across time points.

In the spinal cord, similar changes were observed. *Ppia* was now ranked above *Mrpl10* and these two were identified as the top two (Table 3). However, the third rank was now attributed to *Actb*. *Hsp60*, which was previously third finds itself in the lower half of the ranking. Again, the top three genes identified exhibit low overall variation in the spinal cord and their profiles exhibit minimal variation across timepoints (Fig 3a, f and i).

This indeed shows that exclusion of highly variable genes potentially improves the performance of the NormFinder method and can help in improving the quality of the results.

#### Ranking changes in GeNorm

The changes observed in GeNorm were more drastic in the cerebellum. The new rankings were almost the inverse of the previous rankings (Table 2 & Table 3) with *Actb*/*Rpl13a* now identified as the most stable genes followed by *Rps26, Mrpl10* and *Ppia*. These results, although astonishing, made sense when interpreted with the raw expression profiles (Fig. 2). In the absence of *Gapdh* and *Tbp*, the algorithm has chosen *Actb*/*Rpl13a* to be the most stable. Notice how the profiles of these two genes are very similar (Fig 2a & 2h). The genes ranked just below also have as similar profile to these two. In effect, the previous GeNorm rankings (Table 2) were based on genes that had profiles similar to *Gapdh*/*Tbp* (U shaped curve) whereas the new rankings are based on genes with the exact inverted profile exhibited by *Actb* and *Rpl13a*. This change in ranking brings to evidence the possible biases that can occur when using GeNorm in a heterogeneously correlated data set.

In the spinal cord, however, the changes were not so drastic. The top two genes *Mrpl10*/*Rpl13a* remained the same (Table 2 & Table 3). Nevertheless, the genes that were ranked below showed some noticeable change. *Actb* and *Ppia* are now ranked above *Hsp60*. In the spinal cord, most of the genes are correlated (Fig. 4b) and their profiles look alike except *Rps26* (Fig. 3j). Therefore, the top two genes did not change even after the exclusion of the highly variable genes. However, in the absence of these genes, the profile of *Actb* (Fig. 3a) now looks more like *Mrpl10*/*Rpl13a* than *Ppia* or *Hsp60* (Fig. 3b, 3f, 3h & 3i). This is a possible reason for the change in ranks observed.

These results taken together yet again show that GeNorm results can prove to be highly biased depending on the correlation observed in the dataset and the expression agreement between candidate genes.

#### Ranking Changes in Pairwise ΔCt method

In the cerebellum, the changes observed are very similar to the changes observed in the NormFinder algorithm (Table 3). *Mrpl10* and *Ppia* are still ranked as the top two. However, *Actb* and *Rpl13a* are now ranked above *Sdha, Pgk1* and *Hsp60*. Indeed, the removal of *Gapdh* and *Tbp* decreased the difference in Cq values between *Actb* and other genes thereby leading to lower Mean SD (higher ranks). The same is true for *Rpl13a*. Similarly, genes that resembled *Gapdh* and *Tbp* were now pushed to the bottom as their profiles look much different from the top ranked reference genes. In other words, the difference between their Cq values and the others have increased across all samples. Interestingly, the ranking of the Pairwise ΔCt method and NormFinder seem to concur a lot both before and after exclusion of highly variable genes (Table 2 & Table 3).

However, in the spinal cord, which shows high extent of homogenous correlation, the changes are rather significant. The top two genes now are *Mrpl10* and *Ppia*. This is followed by *Actb*. *Hsp60* which was ranked 1^st^ before exclusion (Table 2) is now close to the bottom of the ranking along with *Rps26* (Table 3). The reason for this change is the same as stated for the cerebellum. Removal of genes with high overall variation results in genes that had a similar profile to the genes removed being classed lower. However, this also depends on the similarity of the rest of the genes with the profile of the new highly ranked genes. This is a significant hurdle as the ranking of genes becomes inter dependent even after the removal of highly variable genes. In the spinal cord, the ranking of this method after exclusion concurs with NormFinder (Table 3), that was previously not the case (Table 2).

## Discussion

As each method analysed in the study has its own advantages and drawbacks, using any of these methods alone would not be enough to have bias-free results. Neither would averaging the ranks of genes determined from all these methods. Hence, there is a need to devise an integrated approach that is based on the suitability of the method in a longitudinal setting.

Among the different methods tested in this developmental study, we find that the methods that rely on pairwise variation (GeNorm and Pairwise ΔCt method) are ill suited for reference gene validation. However, it should be noted that the predictions of the Pairwise ΔCt method did improve after the exclusion of highly variable genes. Nevertheless, the ranking inter-dependency does pose a major problem for a longitudinal study.

Regarding the other two methods, the major drawback of the CV analysis is that it does not factor in the variation between groups but can however determine the absolute overall variation. The major drawback of the NormFinder method is that the stability ranking of the method is influenced by genes with high overall variation. It can however calculate stability based on both intergroup and intragroup variation. Hence, using these two methods in tandem would negate their respective drawbacks. It goes without saying that visual representation of the raw expression profiles is also an additional tool that can be used to validate the findings of these two methods.

Taking all these factors into account, we propose an integrated approach that uses CV analysis, visual representation (with ANOVA) and NormFinder to assess the best set of reference genes for a longitudinal study (Fig. 5). This method also recapitulates the method used in this study.

**Figure 5:**
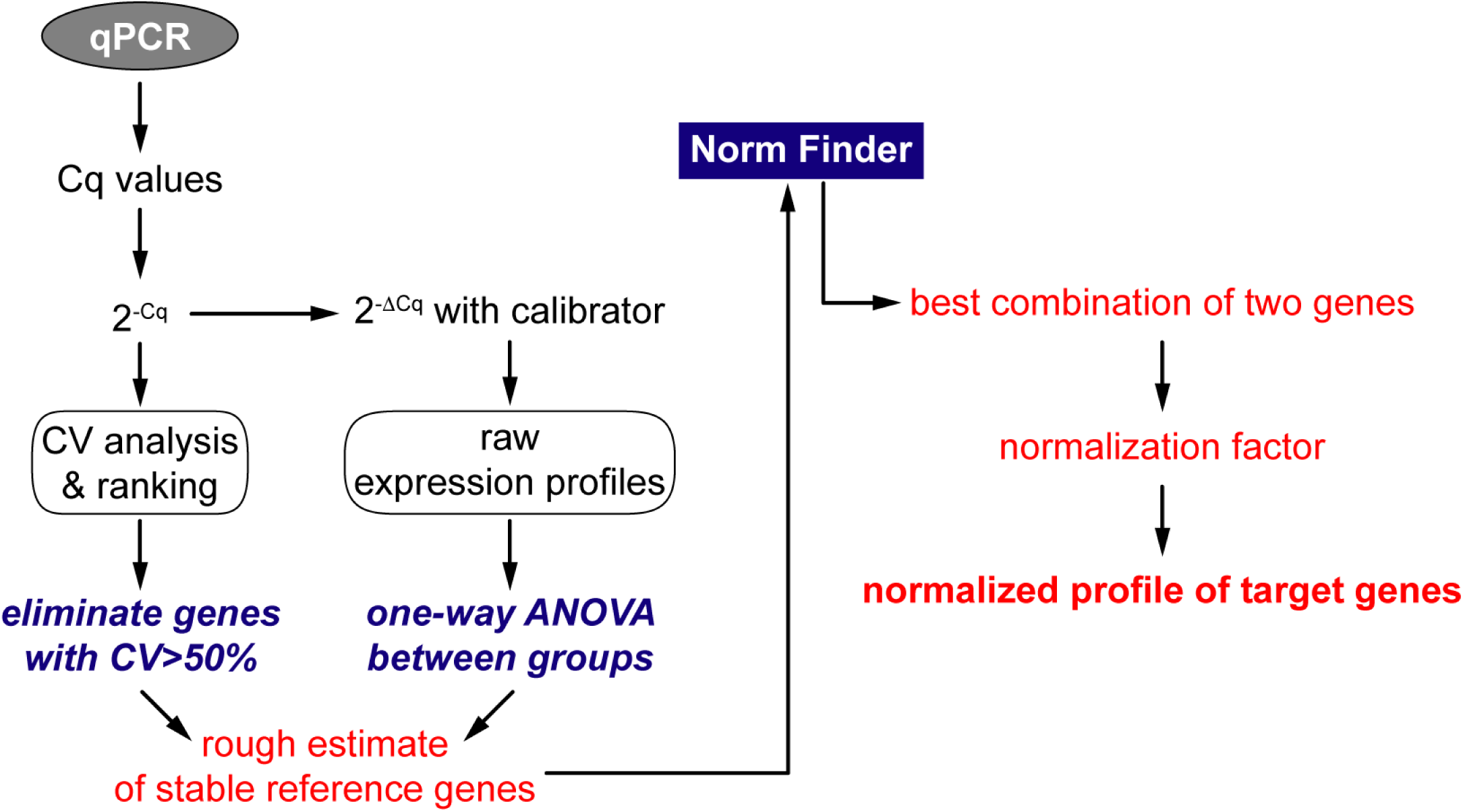
Integrated approach to determine the best reference genes in a longitudinal study. The proposed integrated approach starts by linearizing the Cq values. CV analysis is performed on the linearized values. Genes with CV>50% are omitted from further analysis. Parallelly, using the experimental calibrator, the raw expression profiles are visualised. One-way ANOVA is the applied to check for variation across groups. These two results give a rough estimate of stable references. Then the rest of the genes are subjected to the NormFinder algorithm which determines the two best reference genes which are then used for normalizing target genes.

Once the Cq values of all samples and genes are obtained from qPCR, they are linearized, and CV analysis is performed. Parallelly, using the experimental calibrator, the raw experimental profiles of all candidate genes are plotted as fold changes (2^-ΔCq^). This is followed by One-way ANOVA to assess the variation among all groups. It should be noted that ANOVA is used just to assess if there is significant statistical variation among the means of the groups. It does not assess, by any means, the extent of this variation. Thus, visual representation along with the results of CV analysis and ANOVA can help us obtain a rough estimate of the most stable reference genes. At this stage, genes that exhibit CV>50% are removed. The rest of the genes are then subjected to the NormFinder algorithm. The algorithm then ranks the genes based on intergroup and intragroup variation. It also detects the best two genes that can be used for normalization with a grouped stability value. These two genes are then used to calculate a normalizing factor to normalize all target genes.

In our study, the integrated approach detected *Mrpl10* and *Ppia* as the two best genes in both tissues with a grouped stability of S=0.17 in the cerebellum and S=0.11 in the spinal cord. Nevertheless, we cannot be sure if the same genes would be the best references if another time point was added or if one of the time points studied here is removed. This limitation indeed exists for all reference gene validation studies. In other words, the best reference gene(s) identified by any study using any method is only suitable for that experimental setting and cannot be extrapolated.

Apart from the parameters that have been identified in this study that contribute to the suitability of a method, certain other factors are to be considered while constructing a validation study. Indeed, simple factors such as the total sample size and number of candidate reference genes play an important role on the suitability of a statistical method (22). Similar issues have been detailed and standard guidelines have been established to facilitate standardization and reproducibility of qPCR assays (23).

A notable limitation of this study is the choice of the candidate reference genes. The candidates were chosen as they have been conventionally used for normalization of differential expression in the CNS. This means that we could have potentially excluded unknown and undiscovered stable references. A more thorough approach would be to choose candidate genes from high-throughput data to avoid this inherent bias (24–26).

Furthermore, the commonly used methods described in this study assess stability based on the expression variation of reference genes alone. None of these methods consider the eventual variations that would be present in the target genes tested. Indeed, a reference gene that faithfully mimics the variation observed in the target gene can potentially help in identifying variations caused by the sample and other experimental errors. Including this crucial parameter could indeed increase the reliability of a validation method.

Finally, the integrated approach proposed in this study is only applicable for a longitudinal setting. How these methods would fare in cross sectional studies remains to be explored. However, the approach proposed in this study addresses an important observation that has been constantly made but systematically overlooked in validation studies. Our integrated approach, in effect, is contradictory to previously held notions that all statistical approaches are applicable to any experimental setting and that a minimum of three different statistical approaches are required for robust analyses (27). We remark that these notions can be detrimental to the primary motive of a validation study and it is advisable to devise integrated approaches based on suitability.

## Methods

### Animal Tissue Samples

C57Bl6/J mice at 4 postnatal time points - P5, P10, P15 and P23 were chosen. For each time point, between 5 to 7 animals were sacrificed and their spinal cord and cerebellum were isolated for further experiments. All aspects of animal care and animal experimentation were performed in accordance with the relevant guidelines and regulations of INSERM, Université Paris Descartes and approved by the French National Committee of Animal experimentation and ethics (Ref n°: 2016092216181520).

### Total RNA isolation

Tissues were dissected out from mice at the stipulated time points, immediately frozen in liquid nitrogen and stored at −80°C. Samples were later thawed in 1mL of TRIzol reagent (Ambion Life Technologies 15596018) on ice followed by homogenization in a bead mill homogenizer (RETSCH MM300) with 5mm RNase free stainless-steel beads. Homogenization was carried out for 4 minutes at 20Hz (2 x 2 minutes) with a 3-minute pause in between when the samples were placed on ice to cool down. Next, RNA was extracted from the homogenate using the manufacturer’s instructions with slight modifications. Briefly, 100% Ethanol was substituted for Isopropanol to reduce the precipitation of salts. Also, RNA precipitation was carried out overnight at −20°C in the presence of glycogen. The following day, precipitated RNA was pelleted by centrifugation and washed at least 3 times with 70% Ethanol to eliminate any residual contamination. Tubes were then spin dried in vacuum for 10 minutes and RNA was resuspended in 30μL of RNase Free water (Ambion AM9937). RNA was then stored at −80°C till RT-PCR.

### RNA Quality, Integrity and Assay

RNA quantity was assayed using UV spectrophotometry on Nanodrop One (Thermo Scientific). Optical density absorption ratios A260/A280 & A260/A230 of the cerebellum samples were 1.86(±0.03 SD) and 2.29 (±0.07 SD) respectively. The corresponding values for the spinal cord samples were 1.95 (±0.03 SD) and 2.08 (±0.32 SD) respectively. Furthermore, RNA integrity was verified using denaturing formaldehyde agarose gel electrophoresis. All samples showed intact bands for 28S and 18S rRNA and were subsequently used for RTqPCR.

### RTqPCR

500ng of Total RNA was first subjected to DNase digestion (Promega M6101) at 37°C for 30 minutes to eliminate contaminating genomic DNA. Next, DNase activity was stopped using DNase Stop Solution (Promega M199A) and RNA was reverse transcribed with Random Primers (Promega C1181) and MMLV Reverse Transcriptase (Sigma M1302) according to prescribed protocols. Quantitative Real time PCR (qPCR) was performed using Takyon ROX SYBR 2X MasterMix (Eurogentec UF-RSMT-B0701) as a fluorescent detection dye. All reactions were carried out in a final volume of 7μl in 384 well plates with 300 nM gene specific primers, around 7ng of cDNA (at 100% RT efficiency) and 1X SYBR Master Mix in each well. Each reaction was performed in triplicates. All qPCR experiments were performed on BioRad CFX384.

### Primer Design

All primers used in the study were designed using the Primer 3 plus software (https://primer3plus.com/cgi-bin/dev/primer3plus.cgi). Splice variants and the protein coding sequence of the genes were identified using the Ensembl database (www.ensembl.org). Constitutively expressed exons among all splice variants were then identified using the ExonMine database (www.imm.fm.ul.pt/exonmine/). Primer sequences that generated amplicons spanning two constitutively expressed exons were then designed using the Primer 3 plus software. For detailed information on Primer sequences refer to Table 1.

### Amplification Efficiencies

The amplification efficiencies of primers were calculated using serial dilution of cDNA molecules. Briefly, cDNA preparations from Cerebella and Spinal cords from different age groups of mice were pooled and serially diluted three times by a factor of 10. qPCR was then performed using these dilutions and the results were plotted as a standard curve against the respective concentration of cDNA. Amplification efficiency (E) was calculated by linear regression of standard curves using the following equation

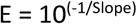

Primer pairs that exhibited an Amplification Efficiency (E) of 1.93 to 2.05 and an R^2^ value (Determination Coefficient) of 0.98 or above were chosen for this study.

### Data analysis & visualization

qPCR readouts were analyzed in Precision Melt Analysis Software v1.2 and the Cq data was exported to Microsoft Excel for further calculations. Differential expression was calculated using the 2^-ΔΔCt^ method (28,29) and data was visualized using Graph Pad Prism v7.0.

### Statistical Analysis

Statistical analysis for reference gene validation was performed using R software packages and Microsoft Excel. GeNorm (18) and Normfinder (19) analysis were performed using published script in R package(30). Co-efficient of Variance Analysis and the pairwise ΔCt analysis was performed in Microsoft Excel. To assess statistical difference in RNA quantities between groups, One-way ANOVA was performed in Graph Pad Prism v7.0. Pearson’s correlation matrix of reference genes and the heat maps were also generated in Graph Pad Prism v7.0.

Author Contributions
V.K.S. and N.K.S. contributed equally to the article. They conceived the study and performed the experiments. V.K.S. analysed the data and wrote the manuscript. C.M. edited and proof-read the manuscript. J.G. prepared the figures. V.K.S. and J.G. revised the article for submission.

## Acknowledgements

The authors would like to thank Paris Descartes University and INSERM. V.K.S. and N.K.S. received their PhD fellowships from the French Ministry of Research and Innovation. The authors would like to acknowledge Céline Becker for the breeding and management of the animals used in the study.

