## Supplementary Information for "Optimal use of statistical methods to validate reference gene stability in longitudinal studies"

**Venkat Krishnan Sundaram^1*^, Nirmal Kumar Sampathkumar^1^, Charbel Massaad^1^, Julien Grenier^1^**

^1^INSERM UMR-S1124, Paris Descartes University, Faculty of Basic and Biomedical Sciences - 45 Rue des Saints Pères, 75006 Paris, France

*Corresponding Author

**Venkat Krishnan Sundaram**

ORCID ID: <https://orcid.org/0000-0001-6024-695>

**SUPPLEMENTARY INFORMATION**

**Supplementary table S1**

**CV analysis calculation: Example**

|  | **CV analysis ACTB** |  |
| --- | --- | --- |
| **Sample** | **ACTB Cq** | **2^-Cq^** |
| **P5_1** | 16.73 | 9.2E-06 |
| **P5_2** | 16.43 | 1.1E-05 |
| **P5_3** | 16.67 | 9.6E-06 |
| **P5_4** | 16.63 | 9.8E-06 |
| **P5_5** | 17.68 | 4.8E-06 |
| **P10_1** | 16.22 | 1.3E-05 |
| **P10_2** | 16.81 | 8.7E-06 |
| **P10_3** | 16.69 | 9.5E-06 |
| **P10_4** | 15.97 | 1.6E-05 |
| **P10_5** | 16.60 | 1.0E-05 |
| **P10_6** | 16.77 | 9.0E-06 |
| **P15_1** | 16.76 | 9.0E-06 |
| **P15_2** | 16.75 | 9.1E-06 |
| **P15_3** | 16.94 | 8.0E-06 |
| **P15_4** | 16.99 | 7.7E-06 |
| **P15_5** | 16.76 | 9.0E-06 |
| **P15_6** | 16.91 | 8.1E-06 |
| **P23_1** | 17.69 | 4.7E-06 |
| **P23_2** | 17.32 | 6.1E-06 |
| **P23_3** | 17.39 | 5.8E-06 |
| **P23_4** | 17.58 | 5.1E-06 |
| **P23_5** | 17.72 | 4.6E-06 |
| **P23_6** | 17.24 | 6.4E-06 |
|  | **Mean 2^-Cq^** | **8.4E-06** |
|  | **SD 2^-Cq^** | **2.7E-06** |
|  | **CV (SD/Mean) x 100** | **31.86%** |

The Cq values of all samples are linearized. The CV is then calculated as the ratio of the SD to the mean of the linearized Cq values. It is expressed as a percentage.

A lower CV value would mean lower variation across samples and therefore higher stability.

**Supplementary Table S2**

**Pairwise ΔCt calculation: Example**

|  | ACTB | HSP60 | GAPDH | SDHA | TBP | MRPL10 | PGK | RPL13A | PPIA | RPS26 |
| --- | --- | --- | --- | --- | --- | --- | --- | --- | --- | --- |
| ACTB vs all | 0.00 | 3.65 | 0.94 | 4.10 | 7.47 | 6.07 | 4.58 | 1.35 | 0.33 | 4.58 |
|  | 0.00 | 3.52 | 0.94 | 4.08 | 7.29 | 6.25 | 4.54 | NA | -0.60 | 4.66 |
|  | 0.00 | 3.82 | 1.01 | 4.18 | 7.76 | 6.21 | 4.72 | 1.24 | 0.42 | 4.50 |
|  | 0.00 | 3.18 | 0.76 | 3.63 | 6.70 | 5.92 | 4.08 | 1.08 | -0.08 | 3.84 |
|  | 0.00 | 3.48 | 0.79 | 3.48 | 6.75 | 5.87 | 4.34 | 1.01 | 0.00 | 4.23 |
|  | 0.00 | 5.44 | 2.05 | 3.67 | 8.50 | 5.45 | 3.53 | 0.59 | -0.05 | NA |
|  | 0.00 | 5.11 | 2.58 | 3.86 | 8.19 | 5.84 | 4.37 | 0.56 | -0.01 | NA |
|  | 0.00 | 5.90 | 2.63 | 4.99 | 9.05 | 6.27 | 4.69 | 1.60 | 1.23 | 4.32 |
|  | 0.00 | 4.86 | 1.96 | 3.65 | 8.44 | 5.54 | 3.85 | 0.82 | 0.11 | 4.17 |
|  | 0.00 | 4.19 | 1.62 | 3.67 | 7.83 | 5.59 | 3.77 | 0.55 | -1.07 | 4.19 |
|  | 0.00 | 4.91 | 1.69 | 3.43 | 8.06 | 5.54 | 3.93 | 0.64 | -0.09 | 4.07 |
|  | 0.00 | 4.41 | 1.35 | 3.55 | 8.90 | 5.75 | 3.93 | 1.34 | 0.32 | 5.00 |
|  | 0.00 | 3.90 | 1.38 | 2.93 | 8.02 | 5.45 | 3.49 | 1.07 | -0.23 | 4.46 |
|  | 0.00 | 4.49 | 1.52 | 3.10 | 8.25 | 5.61 | 3.22 | 0.77 | -0.30 | 4.24 |
|  | 0.00 | 4.11 | 1.56 | 3.01 | 7.89 | 5.68 | 3.53 | 0.95 | NA | 4.60 |
|  | 0.00 | 3.87 | 1.34 | 3.04 | 7.73 | 5.32 | 3.29 | 0.76 | -0.26 | 4.50 |
|  | 0.00 | 4.58 | 1.68 | 3.15 | 8.40 | 5.47 | NA | 0.76 | -0.13 | 4.53 |
|  | 0.00 | 2.91 | -0.84 | 1.76 | 5.87 | 4.68 | 1.90 | 0.94 | -1.23 | 3.84 |
|  | 0.00 | 3.02 | -0.65 | 1.82 | 5.77 | 4.78 | 2.03 | 1.43 | -0.94 | 3.73 |
|  | 0.00 | 2.85 | -0.63 | 1.79 | 5.67 | 4.98 | 2.16 | 1.25 | -0.98 | 4.49 |
|  | 0.00 | 3.03 | -0.79 | 1.70 | 5.83 | 5.18 | 1.90 | 1.25 | -1.04 | 4.18 |
|  | 0.00 | 2.68 | -0.96 | 1.75 | 5.79 | 4.75 | 1.83 | 0.98 | -1.00 | 3.98 |
|  | 0.00 | 2.74 | -0.88 | 1.61 | 5.79 | 5.04 | 2.31 | 0.96 | -0.31 | 4.08 |
| **SD** | **0.000** | **0.913** | **1.143** | **0.957** | **1.123** | **0.472** | **0.996** | **0.299** | **0.601** | **0.315** |
| **Average SD** | **0.758** |  |  |  |  |  |  |  |  |  |

The difference in Cq values between one gene and the others is first calculated (columns) across all samples Each row indicates a particular sample The gene in question here is ACTB. Therefore, the first column has 0s (difference in Cq of the gene with itself)

Next, the standard deviation of the differences is calculated at the end of each column.

The stability value (Average SD) of ACTB is then calculated as the average of these standard deviations

| Corr. Matrix | ACTB | HSP60 | GAPDH | SDHA | TBP | MRPL10 | PGK | RPL13A | PPIA | RPS26 |
| --- | --- | --- | --- | --- | --- | --- | --- | --- | --- | --- |
| ACTB |  |  |  |  |  |  |  |  |  |  |
| HSP60 | -0.141 |  |  |  |  |  |  |  |  |  |
| GAPDH | -0.495 | 0.731 |  |  |  |  |  |  |  |  |
| SDHA | -0.36 | 0.438 | 0.817 |  |  |  |  |  |  |  |
| TBP | -0.44 | 0.802 | 0.937 | 0.735 |  |  |  |  |  |  |
| MRPL10 | 0.572 | -0.021 | 0.06 | 0.431 | 0.037 |  |  |  |  |  |
| PGK | -0.292 | 0.291 | 0.74 | 0.93 | 0.652 | 0.538 |  |  |  |  |
| RPL13A | 0.884 | -0.331 | -0.521 | -0.225 | -0.462 | 0.705 | -0.128 |  |  |  |
| PPIA | 0.393 | 0.267 | 0.095 | 0.212 | 0.175 | 0.546 | 0.307 | 0.544 |  |  |
| RPS26 | 0.802 | 0.095 | -0.217 | -0.13 | -0.084 | 0.615 | -0.098 | 0.791 | 0.405 |  |
| P Values | ACTB | HSP60 | GAPDH | SDHA | TBP | MRPL10 | PGK | RPL13A | PPIA | RPS26 |
| ACTB |  |  |  |  |  |  |  |  |  |  |
| HSP60 | 5.23E-01 |  |  |  |  |  |  |  |  |  |
| GAPDH | 1.63E-02 | 7.53E-05 |  |  |  |  |  |  |  |  |
| SDHA | 9.12E-02 | 3.66E-02 | 2.01E-06 |  |  |  |  |  |  |  |
| TBP | 3.56E-02 | 4.28E-06 | 4.89E-11 | 6.53E-05 |  |  |  |  |  |  |
| MRPL10 | 4.33E-03 | 9.24E-01 | 7.86E-01 | 4.01E-02 | 8.67E-01 |  |  |  |  |  |
| PGK | 1.88E-01 | 1.89E-01 | 8.32E-05 | 3.80E-10 | 1.01E-03 | 9.77E-03 |  |  |  |  |
| RPL13A | 4.95E-08 | 1.32E-01 | 1.29E-02 | 3.13E-01 | 3.05E-02 | 2.46E-04 | 5.80E-01 |  |  |  |
| PPIA | 7.04E-02 | 2.30E-01 | 6.73E-01 | 3.43E-01 | 4.36E-01 | 8.54E-03 | 1.75E-01 | 1.09E-02 |  |  |
| RPS26 | 1.24E-05 | 6.81E-01 | 3.46E-01 | 5.76E-01 | 7.16E-01 | 2.98E-03 | 6.81E-01 | 3.25E-05 | 7.65E-02 |  |

**Supplementary Table S3 : Pearson’s correlation matrix for the cerebellum**

**Supplementary Table S4 : Pearson’s correlation matrix for the spinal cord**

| Corr Matrix | ACTB | HSP60 | GAPDH | SDHA | TBP | MRPL10 | PGK | RPL13A | PPIA | RPS26 |
| --- | --- | --- | --- | --- | --- | --- | --- | --- | --- | --- |
| ACTB |  |  |  |  |  |  |  |  |  |  |
| HSP60 | 0.723 |  |  |  |  |  |  |  |  |  |
| GAPDH | 0.719 | 0.945 |  |  |  |  |  |  |  |  |
| SDHA | 0.75 | 0.934 | 0.889 |  |  |  |  |  |  |  |
| TBP | 0.734 | 0.919 | 0.953 | 0.872 |  |  |  |  |  |  |
| MRPL10 | 0.81 | 0.9 | 0.815 | 0.943 | 0.83 |  |  |  |  |  |
| PGK | 0.745 | 0.958 | 0.893 | 0.97 | 0.886 | 0.934 |  |  |  |  |
| RPL13A | 0.709 | 0.864 | 0.814 | 0.959 | 0.835 | 0.934 | 0.952 |  |  |  |
| PPIA | 0.643 | 0.829 | 0.803 | 0.686 | 0.864 | 0.681 | 0.726 | 0.609 |  |  |
| RPS26 | -0.007 | -0.089 | -0.041 | -0.334 | 0.043 | -0.291 | -0.278 | -0.413 | 0.356 |  |
| P values | ACTB | HSP60 | GAPDH | SDHA | TBP | MRPL10 | PGK | RPL13A | PPIA | RPS26 |
| ACTB |  |  |  |  |  |  |  |  |  |  |
| HSP60 | 6.48E-05 |  |  |  |  |  |  |  |  |  |
| GAPDH | 7.48E-05 | 3.51E-12 |  |  |  |  |  |  |  |  |
| SDHA | 2.46E-05 | 2.50E-11 | 6.62E-09 |  |  |  |  |  |  |  |
| TBP | 4.41E-05 | 2.42E-10 | 6.57E-13 | 2.80E-08 |  |  |  |  |  |  |
| MRPL10 | 1.62E-06 | 2.24E-09 | 1.25E-06 | 5.38E-12 | 5.33E-07 |  |  |  |  |  |
| PGK | 4.58E-05 | 7.40E-13 | 1.01E-08 | 1.87E-14 | 1.93E-08 | 7.13E-11 |  |  |  |  |
| RPL13A | 1.04E-04 | 5.21E-08 | 1.30E-06 | 1.75E-13 | 3.84E-07 | 2.52E-11 | 2.72E-12 |  |  |  |
| PPIA | 6.98E-04 | 5.57E-07 | 2.31E-06 | 2.17E-04 | 5.53E-08 | 2.48E-04 | 8.66E-05 | 1.57E-03 |  |  |
| RPS26 | 9.74E-01 | 6.81E-01 | 8.49E-01 | 1.11E-01 | 8.43E-01 | 1.68E-01 | 1.99E-01 | 4.48E-02 | 8.81E-02 |  |
